# Structural analysis of different LINC complexes reveals distinct binding modes

**DOI:** 10.1101/2020.06.19.161265

**Authors:** Victor E. Cruz, F. Esra Demircioglu, Thomas U. Schwartz

**Affiliations:** Department of Biology, Massachusetts Institute of Technology, Cambridge, MA, 02139, USA

**Keywords:** SUN proteins, KASH proteins, nucleo-cytoplasmic communication, mechanotransduction, nuclear envelope

## Abstract

Linker of nucleoskeleton and cytoskeleton (LINC) complexes are molecular tethers that span the nuclear envelope (NE) and physically connect the nucleus to the cytoskeleton. They transmit mechanical force across the NE in processes such as nuclear anchorage, nuclear migration, and homologous chromosome pairing during meiosis. LINC complexes are composed of KASH proteins traversing the outer nuclear membrane, and SUN proteins crossing the inner nuclear membrane. Humans have several SUN- and KASH-containing proteins, yet what governs their proper engagement is poorly understood. To investigate this question, we solved high resolution crystal structures of human SUN2 in complex with the KASH-peptides of Nesprin3, Nesprin4, and KASH5. In comparison to the published structures of SUN2-KASH1/2 we observe alternative binding modes for these KASH peptides. While the core interactions between SUN and the C-terminal residues of the KASH peptide are similar in all five complexes, the extended KASH-peptide adopts at least two different conformations. The much-improved resolution allows for a more detailed analysis of other elements critical for KASH interaction, including the KASH-lid and the cation loop, and a possible self-locked state for unbound SUN. In summary, we observe distinct differences between the examined SUN-KASH complexes. These differences may have an important role in regulating the SUN-KASH network.

## Introduction

The nucleus in eukaryotic cells is physically separated from the cytoplasm by a double membrane bilayer known as the nuclear envelope (NE). The outer nuclear membrane (ONM) faces the cytoplasm and is contiguous with the endoplasmic reticulum (ER), while the inner nuclear membrane (INM) borders the nucleoplasm. The two membranes are fused at circular openings where nuclear pore complexes (NPCs) reside [1]. The lumen between the INM and ONM is known as the perinuclear space (PNS), has a thickness of 30-50 nm and is contiguous with the lumen of the ER. Communication between the nucleus and the cytoplasm across the NE is accomplished by two separate molecular entities. On one hand, molecular exchange primarily occurs through the NPC [2,3]. On the other hand, mechanical signaling is mediated through molecular tethers, the Linker of Nucleoskeleton to Cytoskeleton (LINC) complexes [4–8].

The core of LINC complexes resides in the PNS, formed by two proteins. SUN (<u>S</u>ad1 and <u>UN</u>C84) proteins traverse the INM, while KASH (<u>K</u>larsicht, <u>A</u>NC-1 and <u>S</u>yne <u>H</u>omology) proteins are tail anchored at the ONM and constitute the cytoplasmic anchor of LINC complexes. SUN proteins are conserved in all eukaryotes [9,10]. They possess an N-terminal domain that projects into the nucleus followed by a transmembrane helix that spans the INM. The nuclear domain of SUN proteins interacts with lamins and heterochromatin, anchoring SUN at the INM[11]. The perinuclear domain of SUN proteins consists of an extended coiled-coil that precedes the conserved C-terminal SUN domain which binds to KASH proteins[12]. To date, five SUN proteins have been identified in humans. SUN1 and 2 are present in all tissues and possess partially redundant functions. SUN3, 4, and 5 are expressed in testis and have a shorter coiled-coil domain than their SUN1/2 counterparts[13].

The tail anchored KASH proteins project their ~20-30 C-terminal residues, the KASH peptide, into the PNS. Their “PPPX” motif at the very C terminus of the KASH peptide is most distinct, highly conserved, and required for binding to SUN proteins [14–17]. Currently, 6 KASH domain-containing proteins have been identified in humans. These are the nesprins (nuclear envelope spectrin repeats), including Nesprin1-4, as well as KASH5 (CCDC155), and KASH6 (LRMP) [18–20]. Many of the known interactions of KASH proteins in the cytoplasm involve cytoskeletal elements. Different Nesprins bind different such elements, suggesting distinct roles for specific SUN-KASH complexes. This is perhaps best illustrated by human diseases, that can result from mutations in individual KASH proteins [21]. For example, KASH-less forms of Nesprin-1 and Nesprin-2 are associated with neurological and muscular defects such as Emery Dreifuss muscular dystrophy (EDMD) and spinocerebral ataxia[22], while mutants of Nesprin-4 are known to cause deafness[23]. Clearly, a detailed molecular understanding of LINC complexes is necessary to understand their distinct biological functions.

The structures of the human heterohexameric SUN2-KASH1 and SUN2-KASH2 complexes have been determined [15,24] and provide a general illustration of the mode of interaction at the core of the LINC complex. The C-terminal ~20 kDa SUN domain of SUN2 folds into a compact β sandwich, which trimerizes into a trefoil, supported by a preceding three-stranded coiled-coil element. KASH1 or 2 bind at the interface between adjacent SUN domains. A loop, disordered in the apo-structure, folds into a beta hairpin and clamps down the KASH peptide to stabilize the interaction. This element is called the ‘KASH lid’[15]. Since each SUN2 trimer binds 3 KASH peptides at once, the architecture of the complex inherently increasing the interaction strength, distributing expected mechanical force across three discrete sites within the complex. SUN2 also contains a loop structure that coordinates a potassium ion (‘cation loop’), which is essential for KASH binding[15]. Finally, a disulfide bond can be formed between SUN2 and KASH1/2 via highly conserved cysteine residues. This likely enhances LINC complex stability even further[25].

To advance our understanding of SUN-KASH pairing, we sought to address this question using structural and biochemical tools. We solved high-resolution crystal structures of SUN2 in complex with KASH3, KASH4, and KASH5, which complement our previously published SUN2-KASH1 and SUN2-KASH2 complexes. The new structures reveal distinct binding interactions of SUN2 with different KASH peptides. In addition, we solved the structure of unbound, apo-SUN2 at high resolution, suggesting a novel mechanism of inactivating Sun. Taken together, this data feeds into the notion that humans have evolved an elaborate LINC-complex network, with the possibility for regulation on multiple levels.

## Results

### SUN2 binds to all 6 human KASH peptides

To begin our study, we first examined direct binding of human SUN2 to all 6 currently known KASH proteins. We co-expressed SUN2 with the predicted perinuclear KASH peptides of Nesprin-1,-2,-3,-4, CCDC155 (KASH-5), or LRMP (KASH-6) in *E. coli*. Throughout the text we will refer to these peptides as KASH1-6, for simplicity. KASH peptides were N-terminally fused to 6 histidines, followed by a GB1 solubility tag [26], and a human rhinovirus 3C cleavage site. SUN2 was expressed with a cleavable, trimerizing-GCN4 (triGCN4) tag [27,28], but without an affinity tag. SUN2-KASH complexes were first nickel affinity purified, thereby eliminating apo-SUN2. After proteolytically cleaving GB1 from KASH and triGCN4 from SUN2, SUN2-KASH complexes were separated by size exclusion chromatography, with free GB1 eluting as a separate peak. This way, we were able verify stable interaction of SUN2 with all 6 KASH peptides (Fig. 1).

**Fig. 1.**
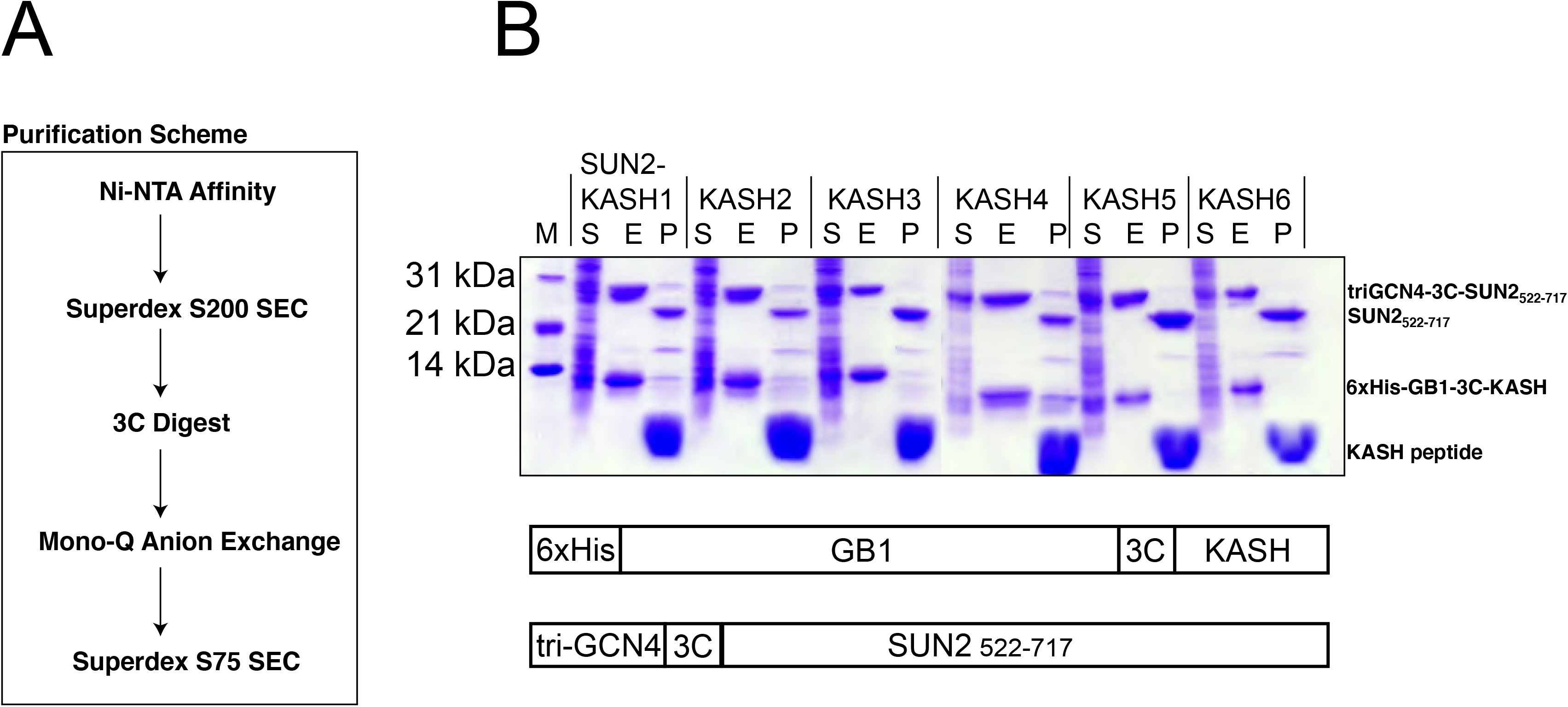
Purification of SUN2-KASH complexes. (**A**) Purification scheme as employed for all SUN2-KASH complex preparations in this study. (**B**) SDS-PAGE analysis of purification steps of SUN2-KASH preparations. S, soluble protein fraction from bacterial lysate. E, Ni-NTA affinity elution fraction. P, pure protein complex after final gel filtration.

### Crystallographic analysis of SUN2 KASH complexes

We next set out to structurally characterize SUN2 in complex with KASH3, KASH4, KASH5, and KASH6 by X-ray crystallography. Using a purification strategy discussed previously [27], our initial attempts yielded poorly diffracting crystals of SUN2-KASH3. After analyzing the crystal packing interactions of these poorly diffracting crystals, we designed SUN2 mutants in the hopes to potentially enhance crystal packing, and hence produce better diffracting crystals[29]. Using this approach, we indeed obtained much better diffracting crystals of SUN2 in complex with KASH3 and KASH5 after introducing 4 point mutations in SUN2: Q534D, T683G, M684R, and A685G. For SUN2-KASH4 crystals, wildtype SUN2 was used. Combined with the new tagging and purification scheme described here we finally obtained well diffracting crystals for apo-SUN2, SUN2-KASH3, SUN2-KASH4, and SUN2-KASH5. (Table 1). All structures pack into crystal lattices related to those observed in previous studies, helping in solving them by molecular replacement[15]. The search model used was SUN2 in complex with KASH1 (PDB code 4DXR), lacking both the KASH lid and KASH peptide. The general features of SUN-KASH engagement are maintained in all complexes (Figure 2). SUN2 forms a trimer, in the apo- and all KASH-bound forms. All KASH peptides bind at the interface between adjacent SUN2 monomers despite sequence variations between KASH peptides. However, we also observe a distinct difference between SUN2 bound to KASH1 or 2 as opposed to KASH3, 4, or 5. While KASH1 and 2 kink toward the periphery of the SUN domain, KASH3, 4, and 5 extend towards the neighboring KASH lid from the adjacent SUN2 monomer (Figure 2). The solvent exposed surface of the KASH lid shows strong conservation that remained unexplained by previous structures of SUN-KASH complexes. The binding mode we observe between SUN2 and KASH3, 4, and 5, reveals that the KASH peptide extends toward these residues, providing a possible explanation for the observed conservation of the solvent exposed face of the KASH lid.

**Table 1.**
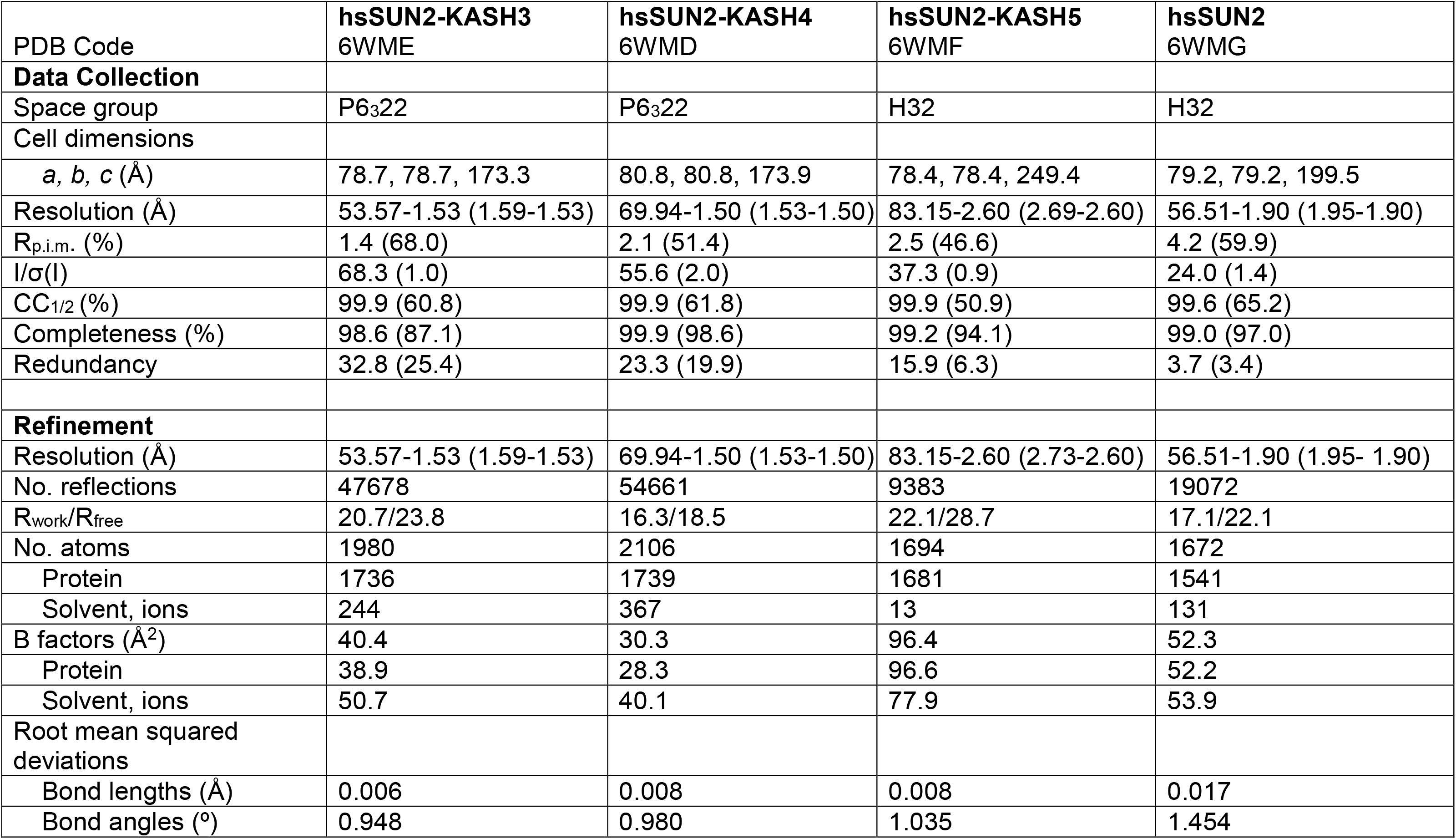

**Fig. 2.**
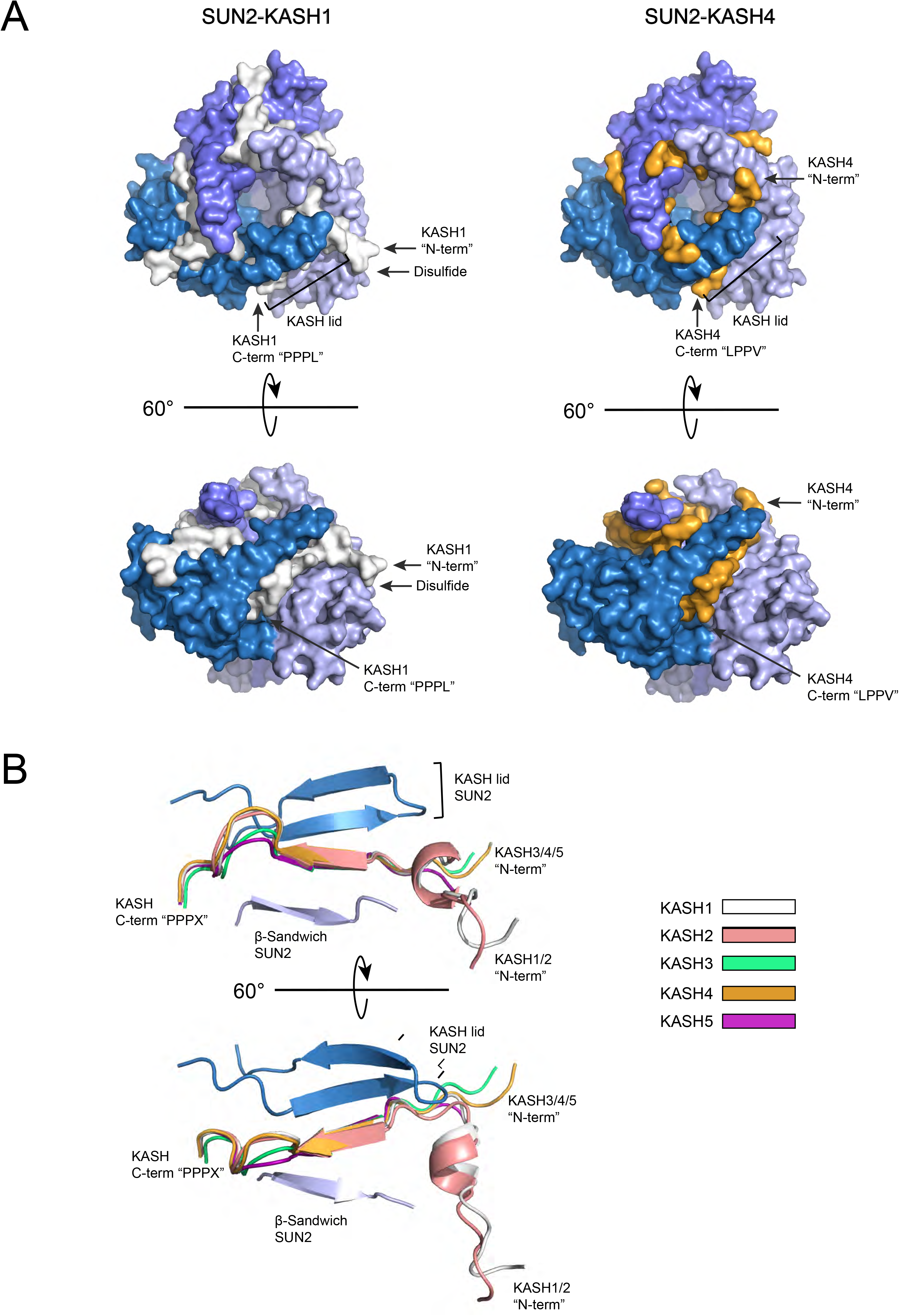
Comparison of Sun-KASH binding modes. (**A**) Space-filling cartoon representation of the two main binding modes, illustrated using the best resolved complex structures for each class (left, SUN2-KASH1, PDB code 4DXR; right, SUN2-KASH4. 6WMD). SUN2 trimer subunits in shades of blue. KASH1 peptide in white, KASH4 peptide in orange. Note the kinked vs. straight conformation of the KASH-peptide. (**B**) Superposition of the five SUN2-bound KASH peptides, including important SUN2 binding elements. The SUN2 conformation is very similar in all five complex structures.

When KASH1, 2, 3, 4, and 5 are aligned, residues between positions 0 (the very C-terminal residue of KASH) to −10 only slightly diverge (Fig. 3). The main differences are start at position −11, which is either a proline or a leucine. When comparing all five KASH peptides side-by-side (Fig. 3, S1), we notice that the peptides with proline in −11 form one class, while the other class has a leucine in the same position. The KASH peptides do not share the same binding surface with SUN2 once they exit from under the KASH lid. Both KASH1 and 2 take a sharp 90° turn at Pro −11 that leads the KASH peptide away from the 3-fold symmetry axis and instead over a neighboring SUN monomer and towards Cys563 where a disulfide bridge is formed. KASH3, 4, and 5 lack Pro −11, and they adopt the alternate binding conformation described here.

**Fig. 3.**
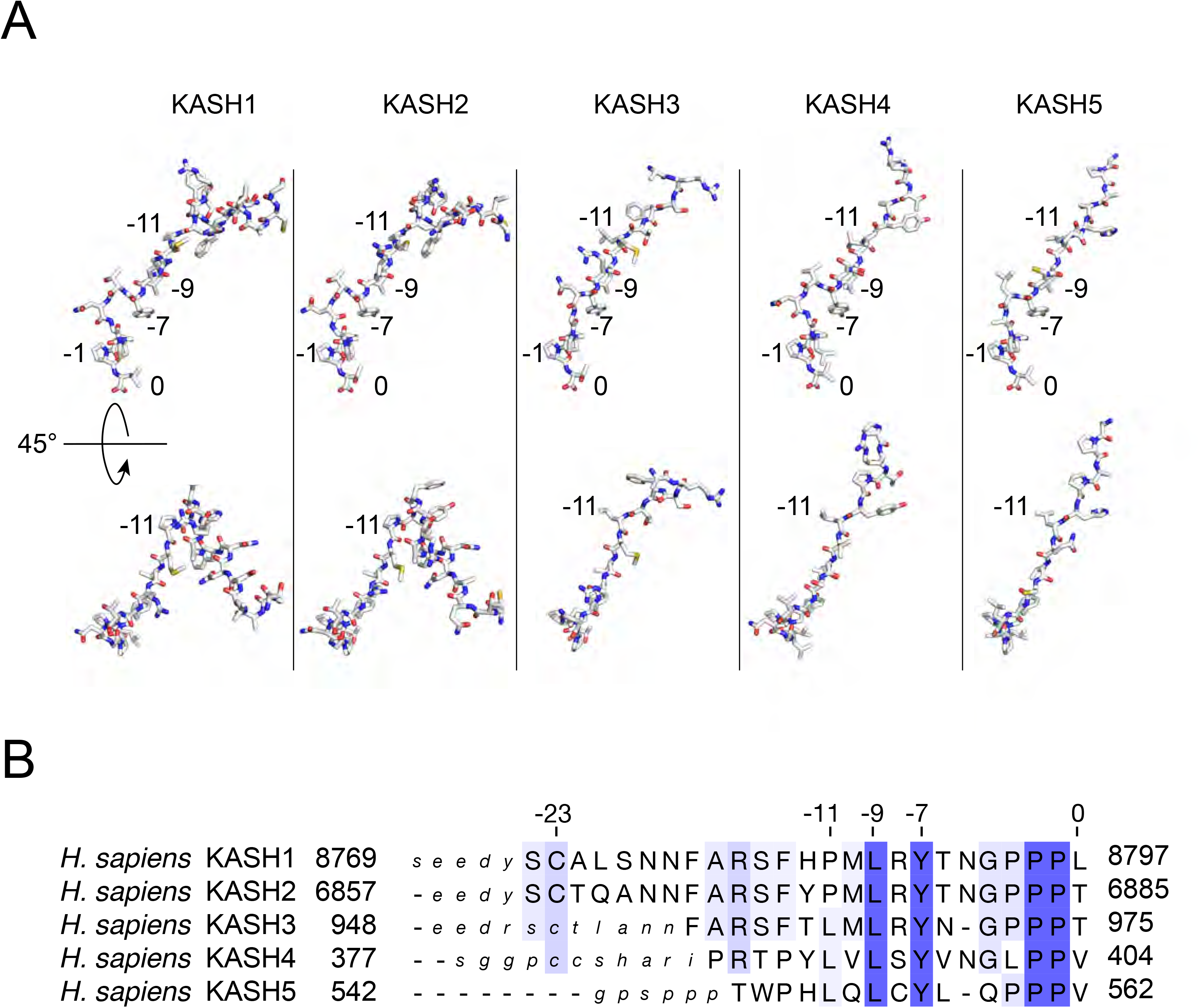
Details of SUN2-bound KASH peptides. (**A**) KASH1-5 are illustrated in stick representation and oriented identically as they are positioned in the SUN2 binding interface. The two views provide detail over the entire KASH sequence. Important positions are numbered, starting at position ‘0’, the very C-terminal residue. Notice position −11, which is where the 90° kink in KASH1 and KASH2 occurs. (**B**) Structure-based sequence alignment of the 5 KASH peptides for which complex structures with SUN2 exist. Sequence conservation indicated in a blue color gradient. Lower case letters represent residues in the crystallized construct that were invisible, i.e. presumably disordered, in the respective crystal structures.

The KASH lid adopts a similar conformation in all five complexes, but it rotates and shifts slightly to accommodate side chains of different sizes present in different KASH peptides. Notably, KASH sequences between position −4 and −6 of different lengths are accommodated (Fig. 2,3) and are partially solvent exposed. KASH3/5 possess two residues here whereas KASH1/2/4 have three. To accommodate the lacking residues at positions −5 and −4, KASH3 and KASH5, respectively, form a shorter loop that still maintains the correct register for the remainder of the KASH peptide. Residues preceding this loop are well aligned between different KASH peptides, and so are residues after this loop. The tyrosine at positions −7 clearly functions as an anchor that is critical in determining the register of the KASH peptide for residues −8 to −11 and facilitates the hydrogen-bonding pattern between the KASH peptide and the KASH lid. The solvent exposed loop between positions −4 to −6 together with the anchoring by Tyr −7 grants flexibility in the sequence and length of residues that can be accommodated between the Tyr −7 and the conserved “PPPX” motif.

### The cation loop of SUN2

We can resolve the electron density of the cation loop in all the structures reported here, but it is especially well ordered in the high-resolution structures of the SUN2-KASH3 and SUN2-KASH4 complexes (Fig. 4). The cation loop of SUN2 in complex with KASH3 is identical to what is observed in the apo form of SUN2, as well as SUN2 bound to KASH1/2/5. In these structures a potassium ion is coordinated at the center of the cation loop by the carbonyl groups of Val 590, Gln 593, Asp 595, Asn 600, Tyr 707, and by a well-ordered water molecule. Surprisingly, the cation loop of SUN2 in complex with KASH4 lacks a cation. Instead of a potassium ion this structure contains two water molecules that coordinate the cation loop interactions. The first water molecule hydrogen bonds with the main chain carbonyl of Val 590, Gln 593, with the main chain amide of Asp 595, and with its neighboring water molecule. The second water coordinates the carbonyl of Asn 600 and of Tyr 707. The main chain carbonyl of Asp 595 now hydrogen bonds with the amine of the Asn 600 side chain and its backbone carbonyl points outside of the cation loop. In the presence of potassium, the side chain of Asp 595 pairs with the amine of the Asn 600 side chain, however, in the absence of potassium the side chain of Asp 595 points away from the cation loop and is no longer as well ordered in our density maps.

**Fig. 4.**
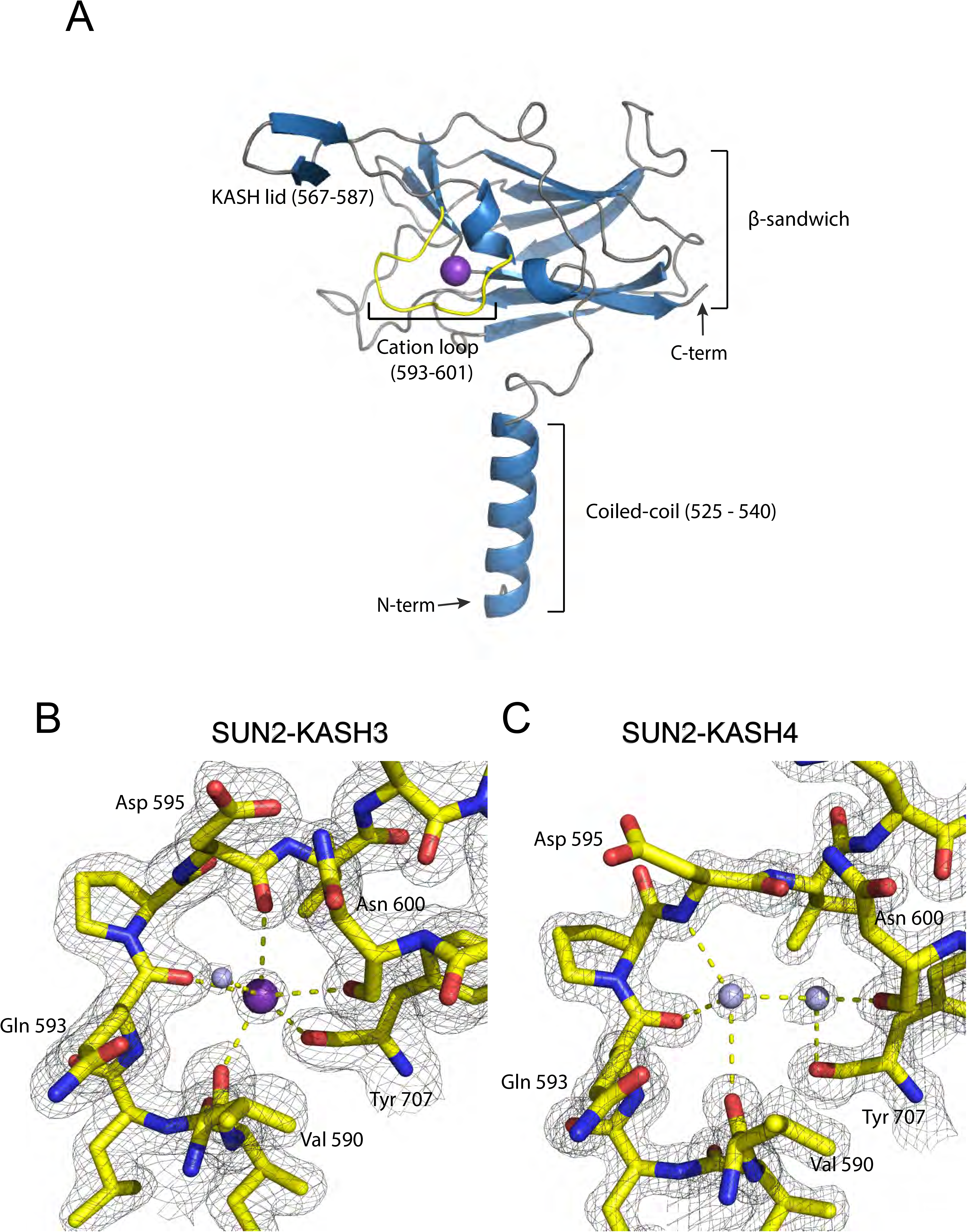
Details of the SUN2 cation loop. (**A**) Overview of a SUN2 monomer, with key structural elements labeled. Purple sphere represents the potassium ion that is bound in most SUN2 structures. (**B**) Detailed view of the canonical potassium coordination in the cation loop, illustrated using the SUN2-KASH3 coordinates. 2F_o_-F_c_ electron density map contoured at 1.5σ. (**C**) The SUN2-KASH4 complex constitutes an exception, as it shows two water molecules positioned in the area that is typically occupied by a potassium ion. 2F_o_-F_c_ electron density map contoured at 1.5σ.

### Internal disulfide of SUN2

SUN proteins contain a pair of universally conserved cysteines (Cys 601 and Cys 705 in SUN2) which form a disulfide bond that presumably stabilizes the cation loop of SUN2. Additionally, this disulfide forms part of the KASH binding pocket that coordinates the conserved “PPPX” motif of KASH (Fig. 5). In the context of KASH1 and KASH2, Cys 601 and Cys 705 of SUN2 form a disulfide. Here we observe that when SUN2 is bound to KASH3, KASH4, or KASH5, these cysteines are reduced. Because of the high quality of our electron density maps the redox state of Cys 601 and Cys 705 is unambiguous.

**Fig. 5.**
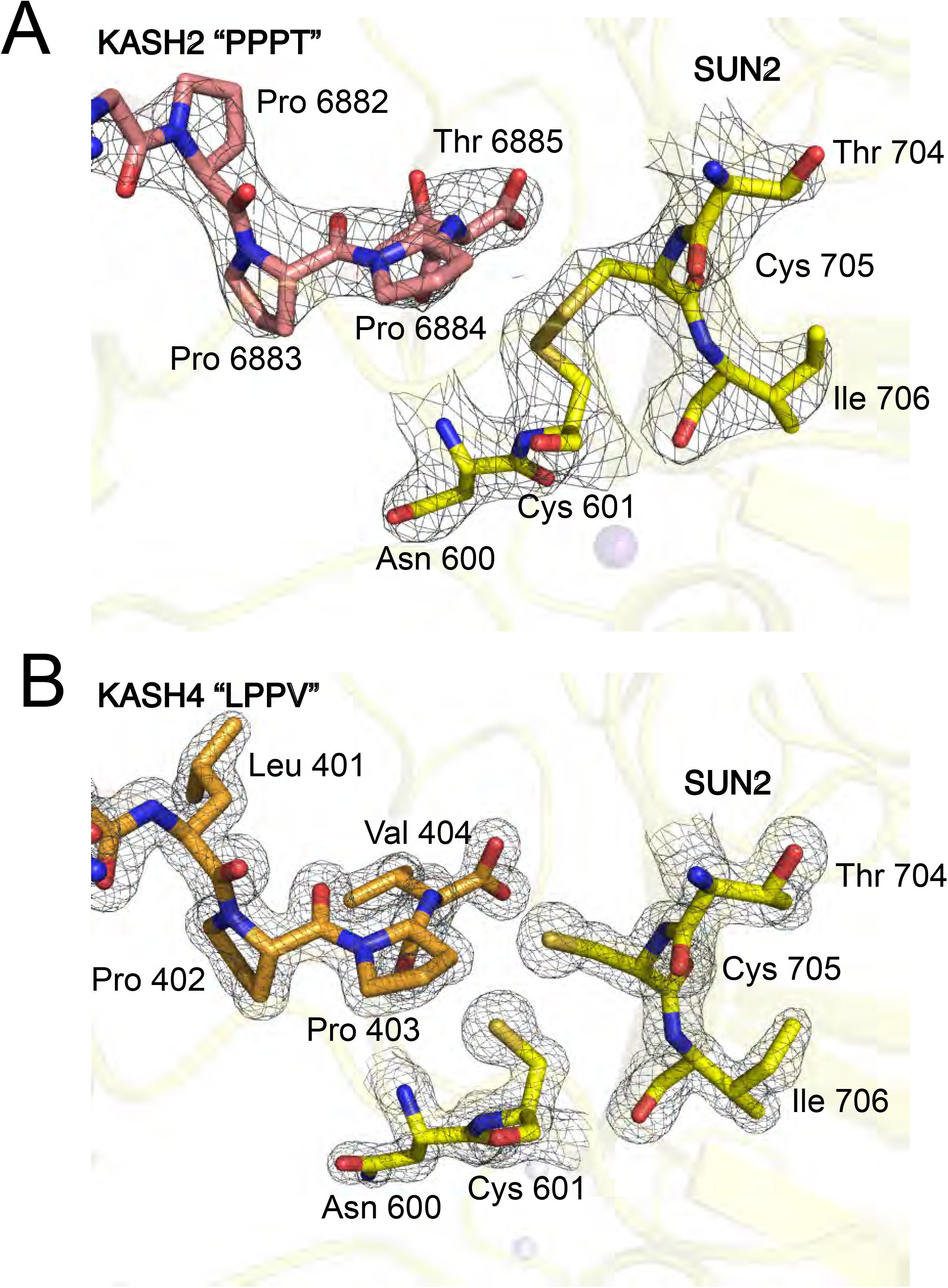
Details of the “PPPX” binding vicinity. (**A**) Detailed view of the SUN2 area in close contact with the C-terminal 3 residues of the KASH peptide. In the KASH1/2 bound structures the cysteine 601/705 pair is best modeled as oxidized, as shown. 2F_o_-F_c_ electron density map contoured at 1.5σ. (**B**) The same detailed view as in (**A**) but showing the KASH4 interaction. Here, the cysteine 601/705 pair is best modeled as reduced. 2F_o_-F_c_ electron density map contoured at 1.5σ.

### The unbound SUN domain may form a self-blocked state

The main motivation behind solving another apo-SUN2 structure was that we extended the construct N-terminally back to residue 500, in the hopes of resolving a longer stretch of the non-canonical trimeric coiled-coil element of SUN2. However, these additional residues are not visible in the structure, suggesting that they are unstructured in the context of this construct. Our new structure superimposes well with the published apo-SUN structures: 0.402 Å RMSD with 4DXT[15] 0.568 Å RMSD with 3UNP[30]. All published and trimeric SUN crystal structures pack similarly, i.e. with the KASH-lid containing, ONM facing side engaging in a head-to-head contact. So far, we concluded that this interaction is non-physiological, primarily on two grounds. First, this conformation would sterically hinder the proper engagement of KASH peptides with the ONM, with the trans-membrane segment of the KASH peptide immediately adjacent to the C-terminal SUN-binding portion. Second, in the KASH-bound form the interaction surface between adjacent trimers is rather small and generally does not pack well. However, in the apo-form this head-to-head crystal interface has a few remarkable features (Fig. 6). In comparison to the apo-form, the KASH-bound head-to-head interface is driven apart substantially, while losing many of the packing contacts. The buried interface in the apo-SUN2 crystal measures 2,380 Å^2^, well in the range of stable interfaces. For KASH-bound interfaces, the buried area measures below 400 Å^2^. In addition, the KASH-lid is dramatically remodeled in the apo-SUN2 form. Overlooked in previous publications, the apo-conformation specifically buries the highly conserved hydrophobic residues Ile 579 and Leu 581 in the head-to-head interface (Fig. 6)[30]. According to PISA analysis, burying these hydrophobics is the main enthalpic contribution driving the interaction. Finally, the well conserved Trp 582 packs tightly against the Glu 552 - Arg 589 salt bridge in the apo-form. The conservation of Ile 579, Leu 581 and Trp 582 cannot be explained by the SUN-KASH interaction, in which these residues are not directly involved. On the other hand, a functional, self-locked apo-state would readily explain the conservation of these residues, including the Glu 552 – Arg 589 salt bridge.

**Fig. 6.**
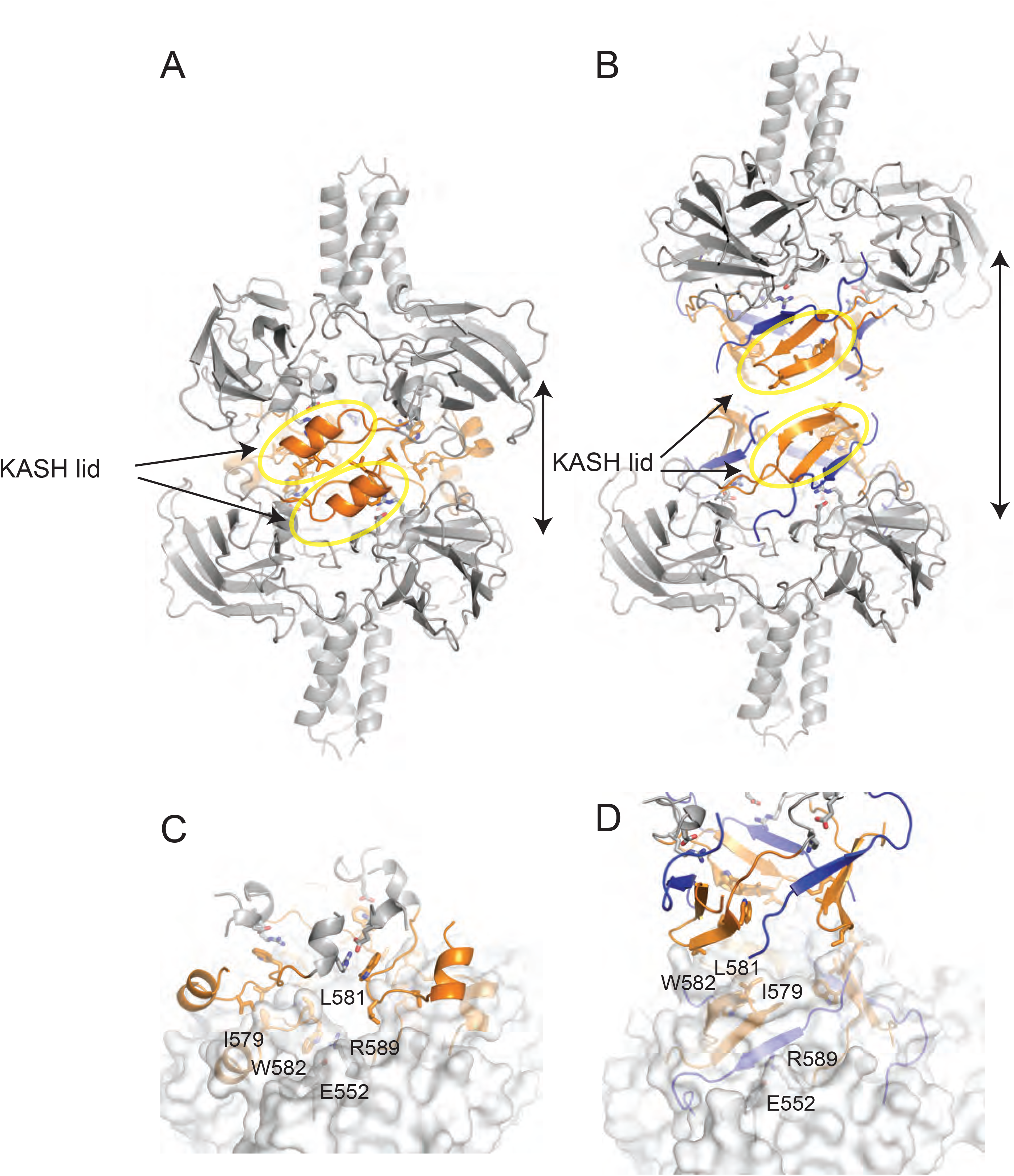
Head-to-head packed apo-SUN2 homotrimers may represent a self-locked state. (**A**) Packing interface in the crystal between two adjacent apo-SUN2 homotrimers. The KASH-lid (orange) is prominently involved in the 2380 Å^2^ interface. (**B**) The same packing interface as in (**A**) but showing the SUN2-KASH4 interaction instead. Notice how the two heterohexamers are much further apart and how the packing interface is significantly smaller. (**C**) an (**D**) are close-ups of the head-to-head interfaces shown in (**A**) an (**B**), respectively. Highly conserved residues involved in the postulated self-locked, apo-SUN2 homohexamer are labeled. The labeled interaction network is repeated 6 times in the apo-SUN2 hexameric interface. In the KASH-bound interface, these residues have no comparable role.

### Mixed occupancy LINC complexes

After confirming that the core interactions between SUN2 and KASH1-5 are similar, we decided to test if a single SUN2 trimer could bind to two different KASH peptides simultaneously and if there is a preference for either binding mode (Figure 7). The same purification scheme shown in Figure 1 was used in this assay, but an additional GFP fused KASH peptide was co-expressed. Since only one KASH type was affinity-tagged, pulling down the other KASH type indicates a mixed LINC complex. This way, we tested if SUN2 could bind to KASH1 and 2 simultaneously. 6xHis-GB1-KASH2 co-purified with both, SUN2 and GFP-KASH1. Similarly, 6xHis-GB1-KASH4 co-purified with SUN2 and GFP-KASH3. These results show that one SUN2-trimer can bind different KASH peptides at once that share the same binding mode. We were also able to purify SUN2 bound to both 6xHis-GB1-KASH2 and GFP-KASH3, simultaneously, showing that SUN2 can bind two KASH peptides that adopt different binding modes. These experiments support the idea that the individual KASH binding sites are independent of one another and that mixed occupancy has to be considered as a means of regulating LINC complexes.

**Fig. 7.**
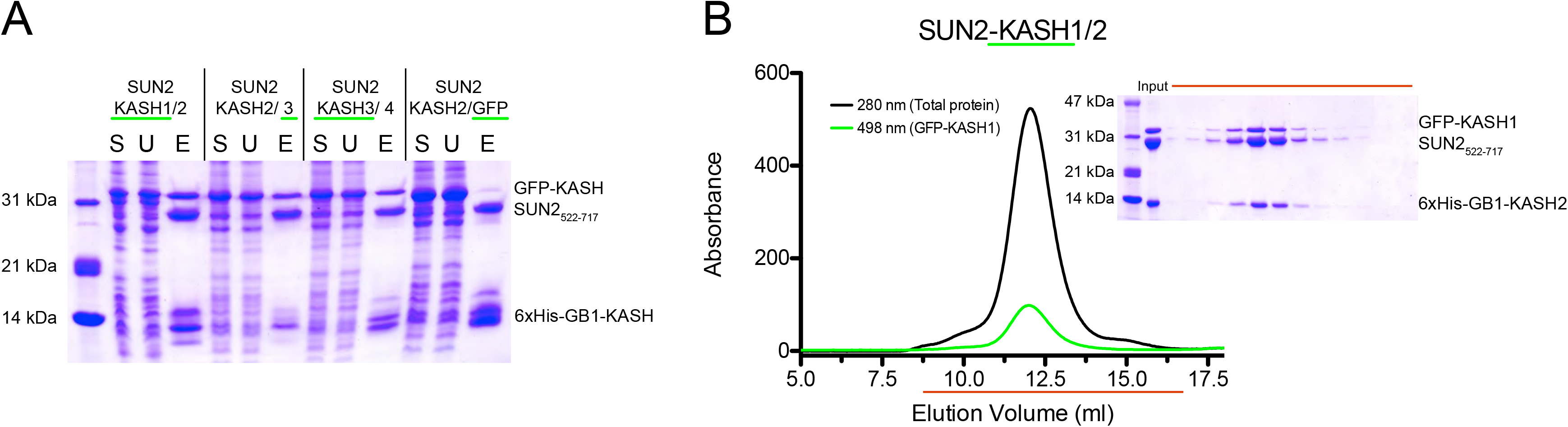
A Sun trimer can bind different KASH peptides simultaneously. (**A**) SUN2_522-717_ was recombinantly co-expressed in bacteria with two KASH peptides, one His-tagged, the other GFP-tagged. The complexes were co-purified using N-NTA affinity, establishing that a SUN2 trimer can simultaneously bind at least two different KASH peptides. The SDS-PAGE shows three KASH combinations and a negative control experiment. S, soluble bacterial protein extract; U, unbound fraction; E, elution fraction from Ni-NTA affinity experiment. (**B**) Representative gel filtration experiment using SUN2_522-717_ / 6xHis-GB1-KASH2/ GFP-KASH1 complex purified via Ni-NTA affinity. Chromatogram shows that GFP-KASH1 coelutes with the main protein peak. SDS-PAGE analysis (inset) of gel filtration fractions (range indicated in red) confirms that all three input proteins coelute, indicating stable complex formation.

## Discussion

Previous work has shown that SUN and KASH interactions occur with promiscuity, which raises the question of how these SUN KASH bridges can be specifically assembled and regulated. Here we show that SUN2 engages with a KASH peptide in at least two distinct binding modes. We can speculate whether a proline or leucine in the −11 position of KASH may be the determining factor to have the peptide engage in a kinked or liner manner, respectively. The argument for such a determination lies within the difference between the binding of KASH1/2 vs. KASH3. While KASH1/2 are kinked, KASH3 is straight. This is remarkable, because these three peptides are very similar in sequence (Fig. 3B) and it is suggestive that the proline to leucine change triggers the conformational change. To be certain, though, this would need to be tested experimentally.

While we now have the structural evidence that SUN2, and most likely the close homolog SUN1 as well, are inherently promiscuous in binding the 6 different KASH peptides, it is not yet clear whether the same will be true for the testis-specific SUN proteins, SUN3, −4, and −5. Those sequences are far enough diverged from SUN2 to preclude a definitive analysis in the absence of structural data.

An outstanding question in the field remains the presumed higher-order assembly of LINC-complexes into 2D clusters [7,31]. Such clusters are thought to be necessary in order for LINC complexes to bear significant mechanical load, i.e. during nuclear migration [25,32]. One easy way to envision cluster formation is that KASH-proteins themselves homo-oligomerize through their cytoplasmic elements. That way, KASH-proteins, together with SUN-trimers, can form higher-order arrays [12]. In this model, one KASH-protein oligomer bridges several SUN-trimers. This is particularly obvious, if the KASH-protein oligomer is not a trimer itself. With our data that formally shows that a single SUN-trimer can bind different KASH-peptides simultaneously, we also have to entertain that higher-order clusters may consist of mixed SUN-KASH complexes. This is an aspect of LINC complex biology that so far has been largely ignored. Another element that has not garnered a lot of attention is the consequence of Nesprins having multiple spectrin repeats, which are protein interaction domains. It is conceivable that molecular condensates, generated through repetitive protein-protein interactions, will have an important role in the molecular function of LINC complexes as well.

We observe subtle differences in the cation loop and the redox state of the close cysteines 601 and 705, depending on which of the five KASH peptides is bound. It remains to be seen whether these differences are biologically meaningful. For example, does the presence of potassium in the cation loop prevent KASH4 interaction, since it is the only structure with water coordination in the cation site? Similarly, is it relevant that the C601-C705 disulfide is reduced in the KASH-3,-4, and-5 bound structures? We do not find any obvious reason, why we see these differences and whether or not they are influenced by the KASH peptides, or whether they may simply be the result of different crystallization conditions. We believe it will be interesting to see more SUN-KASH pairs structurally characterized, at or near atomic resolution, to give a more detailed picture.

Finally, there is an ongoing debate about the regulation of the assembly state of Sun. In order for SUN to bind KASH in a productive and meaningful manner, the cell has to ensure that binding only occurs after SUN has reached the inner nuclear membrane, i.e. traveled from its insertion site within the endoplasmic reticulum, past the circular opening around NPCs, to its destination. If SUN binds KASH before passage to the INM, it will be sterically blocked to pass the NPC, and therefore presumably be non-functional. One way to envision that KASH engagement only happens after SUN reaches its destination is that Sun is kept in a non-binding state before. Studies by the Feng lab suggest that SUN1/2 can adopt a monomeric, KASH-binding incompetent state[33]. In these structural studies the perinuclear coiled-coil element engages with the SUN domain to occlude the KASH binding site. Thus, one can envision that Sun is kept in this inactive form until it reaches the INM. Once there, a yet-to-be-identified signal could switch SUN from inactive to active. We hypothesize that the head-to-head self-locked apo-SUN structure we observe may better reflect a KASH-binding inhibited state. First, the monomeric SUN form described in [33] is dependent on a specific coiled-coil truncation being present. Whether this is representative of the behavior of full-length SUN is not proven, a significant limitation to the study. In contrast, for our self-locked structure, there is no principal concern why it could not be physiologically relevant. In fact, since the self-locking would work by tilting the SUN domains away from the ONM, it would be an elegant and simple solution to prevent KASH binding. Our apo-structure now provides a framework for specifically studying a self-locked state *in vivo.*

In summary, we report near-atomic resolution structures of several SUN2-KASH complexes that shed light on the details of the mode of interaction. Also, we formally show that a Sun trimer can engage with different KASH proteins simultaneously. Much is still to be learned about the nuclear and cytoplasmic anchorage of LINC complexes, as well as functional assembly states, important elements on the path toward a full molecular understanding of mechanotransduction across the NE.

## Materials and Methods

### Plasmid construction, protein expression and purification

DNA sequences containing human SUN2 were cloned into a modified bicistronic ampicillin resistant pETDuet-1 vector (EMD Millipore), superfolder GFP-KASH fusions (sfGFP) were cloned and expressed into a modified ampicillin resistant vector, pET-30b(+) (EMD Biosciences). Each KASH peptide (KASH1-6) was C-terminally fused to 6xHistidine-sfGFP using inverse PCR. For crystallography, 6xHistidine tagged SUN_2522-717_ was cloned into the first multiple cloning site (MCS) and MBP-fused KASH4379-404 was cloned into the second MCS. For SUN2-KASH_3947-975_ and SUN2-KASH_5542-562_ the KASH peptides were C-terminally fused to 6xHistidine-GB1 tags and SUN_2522-717_ remained untagged. All SUN2 and KASH constructs were N-terminally fused with a human rhinovirus 3C protease cleavage site.

Transformed LOBSTR(DE3)-RIL bacterial expression cells[34] were grown at 37 °C to an OD_600_ of 0.6, then shifted to 18 °C and induced with 0.2 M isopropyl β-D-1-thiogalactopyranoside for 16 hours. Cells were harvested by centrifugation at 6000 g, resuspended in lysis buffer (50 mM potassium phosphate, pH 8.0, 400 mM NaCl, 40 mM imidazole) and lysed using an LM20 Microfluidizer Processor (Microfluidics). The lysate was cleared by centrifugation at 10000 g for 25 minutes. The soluble fraction was incubated with Nickel Sepharose 6 Fast Flow beads (GE Healthcare) for 30 minutes at 4 °C in batch. After the beads were washed with lysis buffer, the protein was eluted (10 mM Tris/HCl pH 8.0, 150 mM NaCl, 250 mM imidazole). The protein was further purified by size exclusion chromatography using an S75 or S200 16/60 column (GE Healthcare) equilibrated in running buffer (10 mM Tris/HCl, pH 8.0, 150 mM NaCl, and 0.2 mM EDTA). Protein affinity- and solubility-tags were removed with 3C protease and the protein complexes were separated from fusion tags by a second size exclusion step using an S75 16/60 column, under the same conditions.

### Crystallization

Purified Apo-SUN2_500-717_ was concentrated to 7 mg/ml and crystallized in 14.5% (w/v) polyethylene glycol (PEG) 3350, 100 mM potassium formate pH 8.0, and 200 mM sodium bromide. Rhombohedral crystals appeared after 5 days at 18°C. SUN2-KASH3 complex was concentrated to 5 mg/ml, and crystallized in 16-18% PEG3350, 100 mM ammonium citrate pH 7.0, 100 mM BisTris/HCl pH 6.5, and 10 mM nickel (II) chloride. Large, plate-shaped crystals grew within 2 days at 18°C. SUN2-KASH4 crystallized at 7 mg/ml in 17% PEG3350, 200 mM magnesium chloride, and 100 mM BisTris/HCl pH 6.5. Crystals appeared after 1 day and finished growing within 3 days, at 18°C. SUN2-KASH5 was crystallized at 10 mg/ml, in 14% PEG3000, and 100 mM BisTris/HCl pH 6.5. Crystals appeared and finished growing within 1-12 hours at 4°C. Crystals were cryoprotected in their reservoir solution supplemented with 15% glycerol and flash-frozen in liquid nitrogen. Diffraction data were collected at beamline 24ID-C at Argonne National Laboratories.

### X-ray data collection and structure determination

All data processing was done using programs provided by SBgrid[35]. Data reduction was carried out with HKL2000[36], molecular replacement with PHASER from the phenix suite[37], using apo-SUN2 structure (PDB code 4DXT) that lacked the KASH lid as a search model. The structures were manually built using Coot[38] and refined with phenix.refine. Data collection and refinement statistics are summarized in Table 1. Structure figures were created using PyMOL[39].

### Accession numbers

Coordinates and structure factors have been deposited in the Protein Data Bank under accession codes PDB **6WME** (SUN2-KASH3), **6WMD** (SUN2-KASH4), **6WMF** (SUN2-KASH5), and **6WMG** (apo-SUN2).

## Acknowledgments

This work was supported by NIH grant R01-AR065484 to T.U.S., the NIH Pre-Doctoral Training Grant T32GM007287 (V.E.C.) and is based on research conducted at the Northeastern Collaborative Access Team beamlines, which are supported by NIH grants P30GM124165 and S10OD021527, and DOE contract DE-AC02-06CH11357. We thank the staff at the Advanced Photon Source beamline 24-ID-C, K. Rajashankar in particular, for assistance in data collection and space group analysis.

## Figure legends

Fig. S1. **Details of SUN2-bound KASH peptides, including the SUN2 surface.** (**A**) KASH peptides lined up in the same way as in Fig. 3 but including the SUN2 surface.

