## Supplementary figures and images for "Structural analysis of different LINC complexes reveals distinct binding modes"

### Supplementary Figure S1

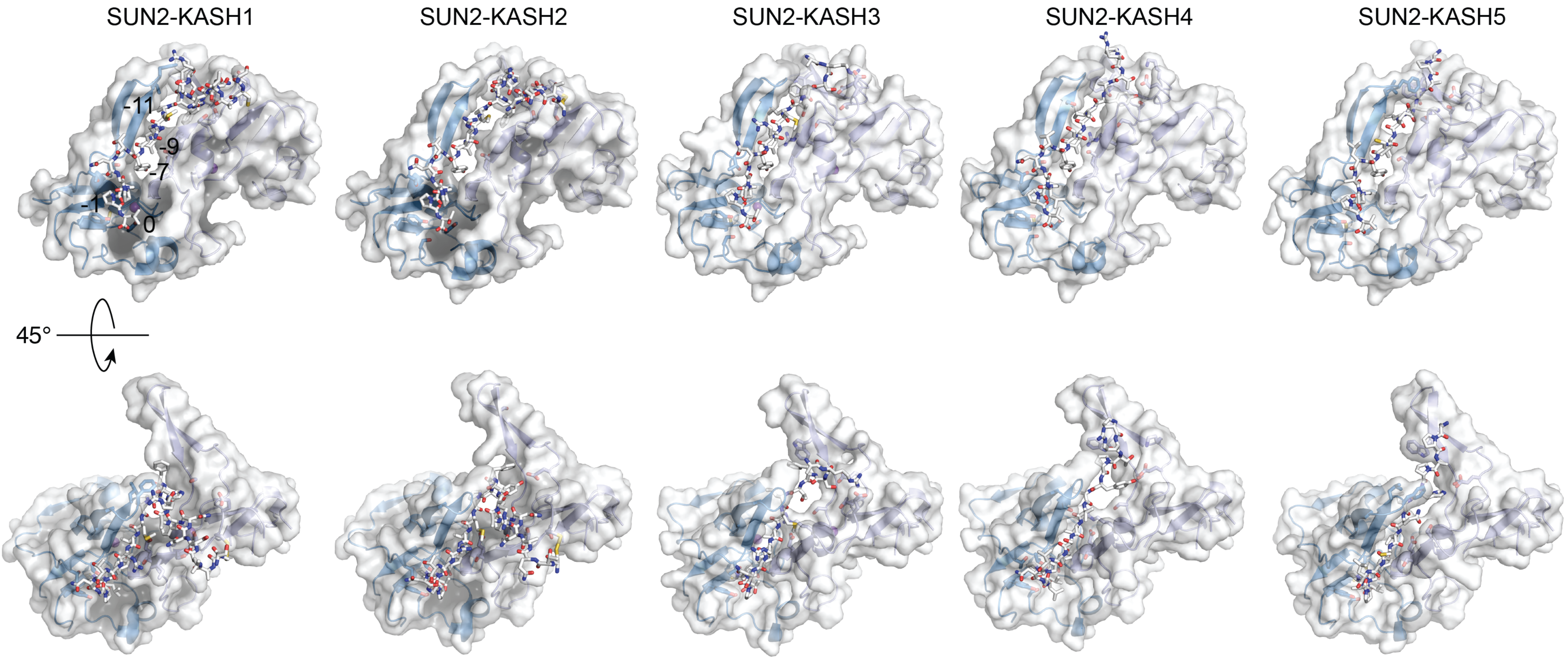
